# Comparing chromatin contact maps at scale: methods and insights

**DOI:** 10.1101/2023.04.04.535480

**Authors:** Laura M. Gunsalus, Evonne McArthur, Ketrin Gjoni, Shuzhen Kuang, Maureen Pittman, John A. Capra, Katherine S. Pollard

## Abstract

Comparing chromatin contact maps is an essential step in quantifying how three-dimensional (3D) genome organization shapes development, evolution, and disease. However, no gold standard exists for comparing contact maps, and even simple methods often disagree. In this study, we propose novel comparison methods and evaluate them alongside existing approaches using genome-wide Hi-C data and 22,500 *in silico* predicted contact maps. We also quantify the robustness of methods to common sources of biological and technical variation, such as boundary size and noise. We find that simple difference-based methods such as mean squared error are suitable for initial screening, but biologically informed methods are necessary to identify why maps diverge and propose specific functional hypotheses. We provide a reference guide, codebase, and benchmark for rapidly comparing chromatin contact maps at scale to enable biological insights into the 3D organization of the genome.

## Introduction

The same genomic locus can adopt different three-dimensional (3D) conformations in different cells, species, and disease states, which can impact gene regulation, cell identity, and replication timing (**Fig. 1A**)^1, 2, 3–7^. Chromosome-conformation capture methods (3C, 4C, 5C, Hi-C, Micro-C)^8–12^ measure how the genome folds across scales, including chromosomal territories, topologically associating domains (TADs), enhancer-promoter loops, and architectural stripes ^10, 13–15^. In recent years, single-cell and deep learning techniques accelerated the study of chromatin conformation across an expanding range of biological contexts ^16–22^.

**Figure 1.**
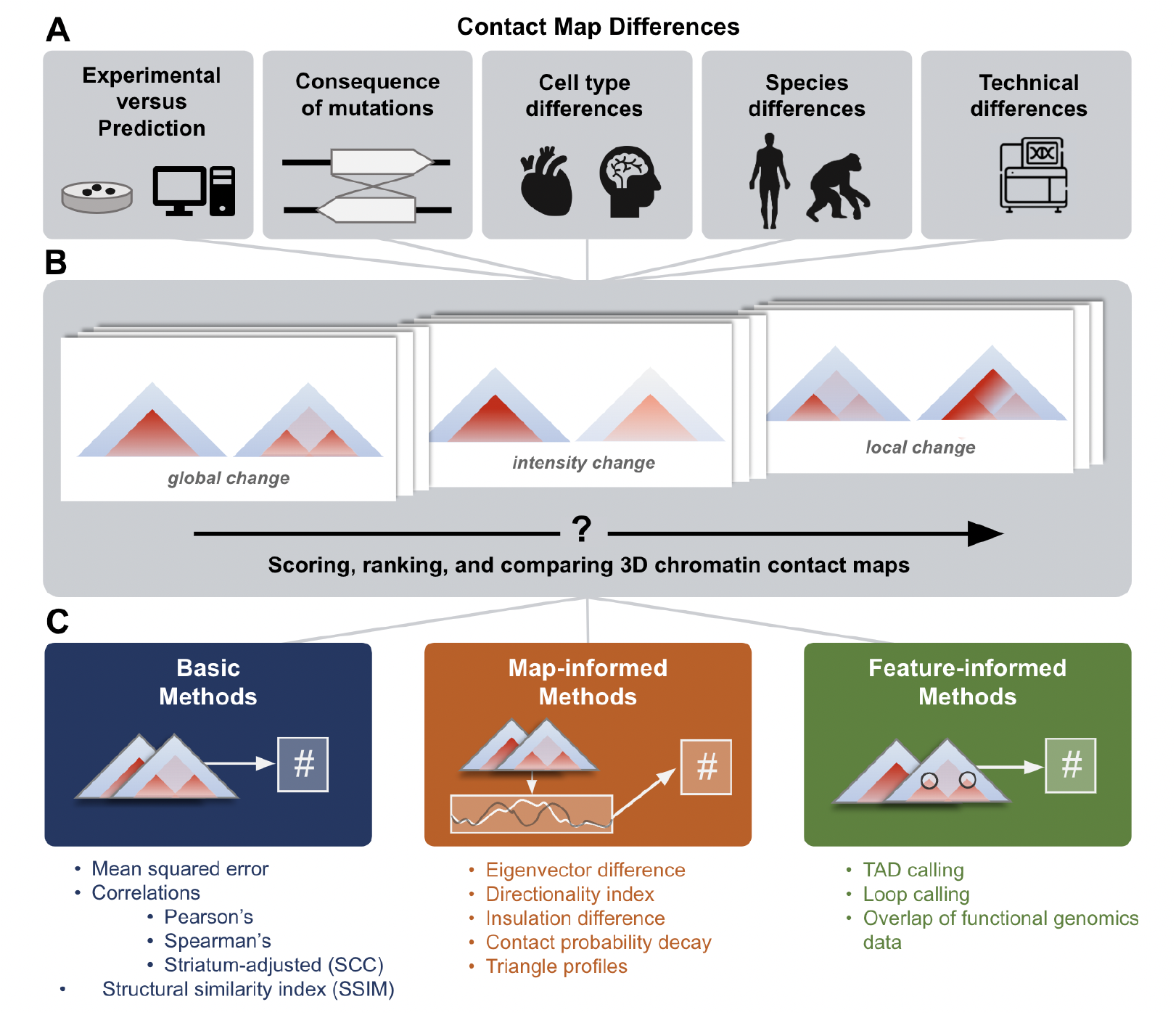
Approaches for comparing 3D chromatin contact maps. (**A**) 3D genome comparisons drive insights into many domains of chromatin biology. Differences observed between maps may reflect consequences of mutations, cell type differences, species differences, or technical biases. (**B**) 3D contact maps exhibit a range of functionally meaningful differences, e.g., in global folding patterns, contact intensity, or small, focal changes to part of the map. (**C**) We define three categories of comparison methods and evaluate 11 representative methods. *Basic methods* (left) compare the contact intensities at each contact bin across two maps with simple measures such as mean squared error or correlations. *Map-informed methods* (middle) transform the 2D contact maps into 1D tracks that describe qualities like the directionality index or insulation score. These tracks were compared to obtain a score. *Feature-informed methods* (right) are designed to identify relevant elements (e.g., from functional genomics data) or structures (e.g., TADs or loops).

There are many ways to compare chromatin conformation maps, but no gold standard exists. Existing approaches rank differences between pairs of maps^6, 7, 23–26^, test reproducibility between replicates and modalities ^7, 23, 24, 27^, identify tissue specific contacts^26^, and highlight differential chromatin interactions^6, 25^. Some scores are designed to identify global differences like boundaries and contact intensities (**Fig. 1B, left and center**), while others target focal changes like enhancer stripes (**Fig. 1B, right**). To rank thousands of loci with diverse folding patterns, one must consider how scoring metrics prioritize different map features and respond to technical artifacts.

Here, we develop a unifying framework to guide strategies for comparing contact maps for new use cases. We introduce three novel methods—eigenvector difference, contact decay probability difference, and triangle track comparison—and benchmark these along with representative methods from the literature to evaluate 11 total approaches (**Fig. 1C**). We quantify how methods differentially rank pairs of contact maps across experimental Hi-C data, 22,500 *in silico* sequence insertions and deletions, and simulated contact maps that capture both biological and technical variation. Our analyses identify when methods diverge and when they are consistent, which methods are redundant or complementary, and where methods commonly fail. The new methods we introduce have relatively high concordance with existing metrics while providing rich information about biological mechanisms. We summarize our recommendations and release a library of open-source code for scoring differences between contact frequency maps to enable scientists to choose and apply the right method for their research question.

## Results

### Diverse strategies for scoring pairs of contact maps

When scoring differences between pairs of contact maps, it is common to apply *basic* methods that consider entire 2D contact matrices (e.g., mean squared error ^206,720^) or *feature-informed* methods that sum differences in specific structures (e.g., loops^28^). These methods represent two extremes. Basic methods are global summary statistics that can overlook small differences that are most biologically interesting. In contrast, feature-informed algorithms specifically target elements such as TADs, stripes, and loops, but are agnostic to overall contact change and may emphasize artifactual differences. As a compromise between these extremes, we extend statistics previously developed to quantify compartments (eigenvectors/PCA^29^), boundaries (directionality index^30^, insulation^31^), and contact decay^9^ in individual maps to instead score differences between pairs of maps. We also propose a new method, called triangle score, which calculates average contact frequencies across all submatrices in a larger contact matrix. These new *map-informed* methods (**Supplemental Text**) transform 2D contact matrices into 1D tracks that capture features relevant to genome folding, and then score them using Spearman’s correlation or mean squared error (MSE). The intermediate 1D track allows for the interpretation of which regions contribute most.

To comprehensively characterize the behavior of the basic, map-informed, and feature-informed scoring approaches, we implemented 11 representative methods in open-source code (**Fig. 1C, Supplementary Table 1, Supplemental Text**): MSE, Spearman’s rank correlation coefficient (ρ), structural similarity (SSIM), stratum-adjusted correlation coefficient (SCC), eigenvector difference, directionality difference, insulation difference, contact probability decay difference, triangle score, the HiCCUPS loop caller^28^, and the cooltools TAD caller^31, 32^. We evaluated how these methods perform across diverse settings. We first applied the methods to Micro-C from human foreskin fibroblasts (HFF) and embryonic stem cells (ESC) to develop biological intuition about the type of map differences each method captures. We then evaluated their performance using a mass screen of *in silico* genetic perturbations. Finally, simulations isolated the effects of specific kinds of technical and biological variation. This three-part benchmark focuses on how methods rank map pairs, rather than the statistical significance of specific differences; stricter or looser significance thresholds can be applied to any score. In sum, we explored and quantified the behavior of scoring methods to learn when they are discordant with each other.

### Beware! Map comparison methods produce discordant rankings

Spearman’s correlation, Pearson’s correlation, and mean squared error are most commonly used to score two maps^28, 33, 34^, as they are computationally efficient and require no feature selection. We compared their behavior using Micro-C contact maps from HFF and ESC cells across all 7,840 1-Mb windows of the human genome (**Methods**). These basic methods prioritized markedly different regions (**Fig. 2**, r^2^ = 0.0002, **Supplemental Fig. 1**)^11^, often for reasons unrelated to underlying biology. For example, a pair of maps with visible structural rearrangements but a low range of contact frequencies was prioritized by correlation, but not by MSE, as the absolute difference between them is small (**Fig. 2A**). Conversely, two maps with similar overall structure but different contact frequency ranges produce a large MSE even though they are very strongly correlated with each other (**Fig. 2D**). These inconsistencies occur because Spearman’s correlation is agnostic to intensity changes, while MSE is sensitive to intensity. Basic methods were not designed to identify specific chromatin features, and therefore may not always be biologically interpretable on their own. They often disagree.

**Figure 2.**
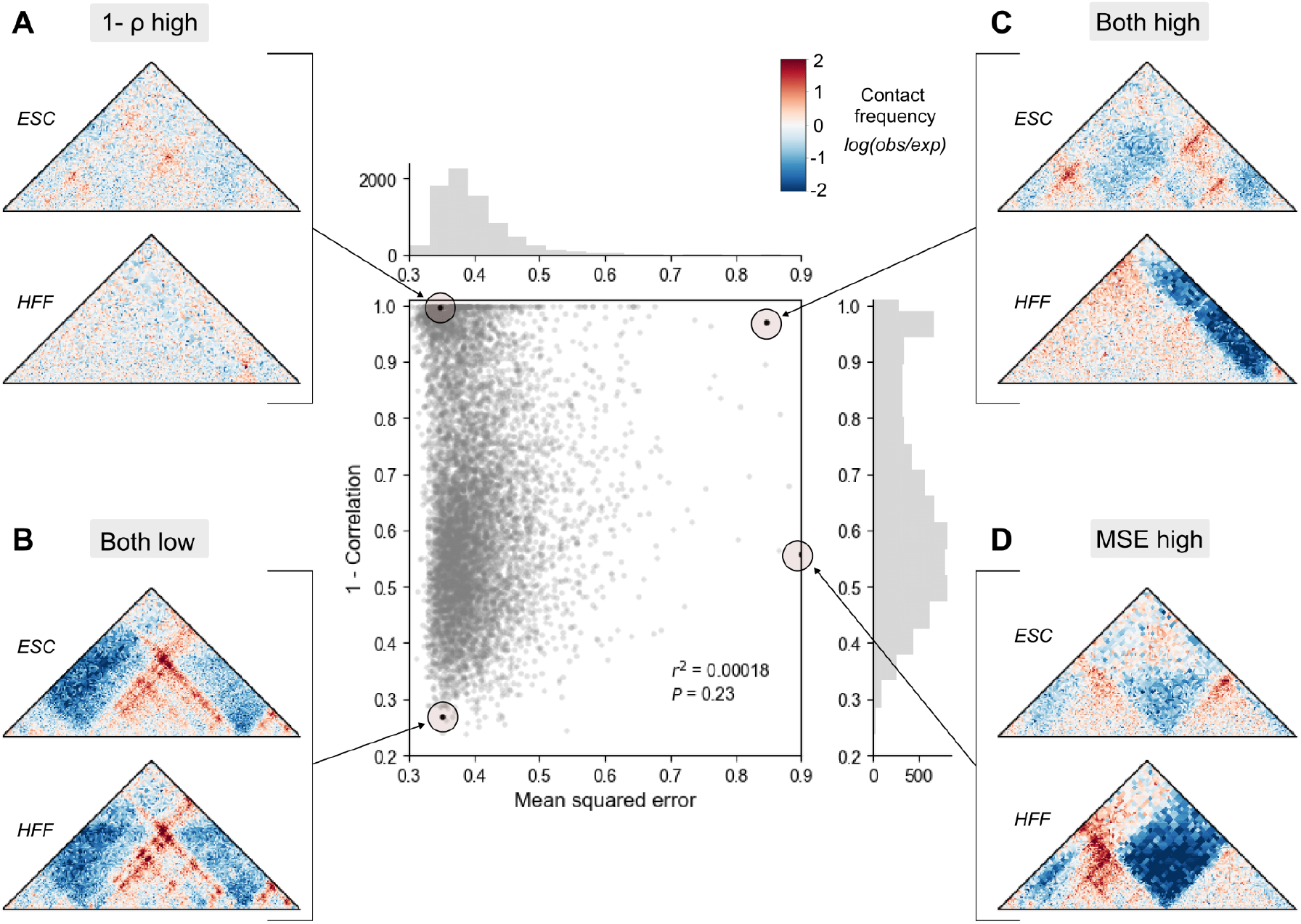
Basic methods to compare contact frequency maps rank map pairs differently. Mean squared error (MSE) and Spearman’s correlation (ρ) were calculated across the genome on experimental contact maps from embryonic stem cell (ESC) and human foreskin fibroblast (HFF) (*n* = 7840). Each point represents a comparison score between a pair of contact maps. We highlight examples where (**A**) only correlation ranks highly, (**B**) both methods agree the maps are similar, (**C**) both methods agree the maps are different, and (**D**) only MSE ranks highly.

### Map-informed methods highlight changes in genome structure

The *map-informed* methods we created or extended have never been benchmarked. To gain intuition about their behavior, we used our comparison across experimental Micro-C maps in HFF and ESCs to evaluate how these methods behave on contact maps containing three common changes linked to disruption in gene regulation: a boundary change, a stripe change, and a loop change (**Fig. 3A i, red boxes**). Triangle score, directionality index, insulation difference, and eigenvector difference all correctly identified large contact changes across the three examples (**Fig. 3A ii-v**). Eigenvector difference in particular showed a strong separation between tracks at the emergence of a new boundary and the strengthening of an existing boundary (**Fig. 3A iii**). Compared to other approaches, directionality index performed best in identifying focal changes, like the loss of loops (**Fig. 3A iv**), while eigenvector difference and insulation difference instead prioritized global changes in contact. Finally, eigenvector difference and contact decay were sensitive to overall contrast difference. We observed a divergence in the contact decay tracks across the first pair where a map gains distal contact **(Fig. 3 vi)**. In sum, the design of these methods highlight different features in the tracks, from overall structural differences and average contact, to sharp changes in contrast.

**Figure 3:**
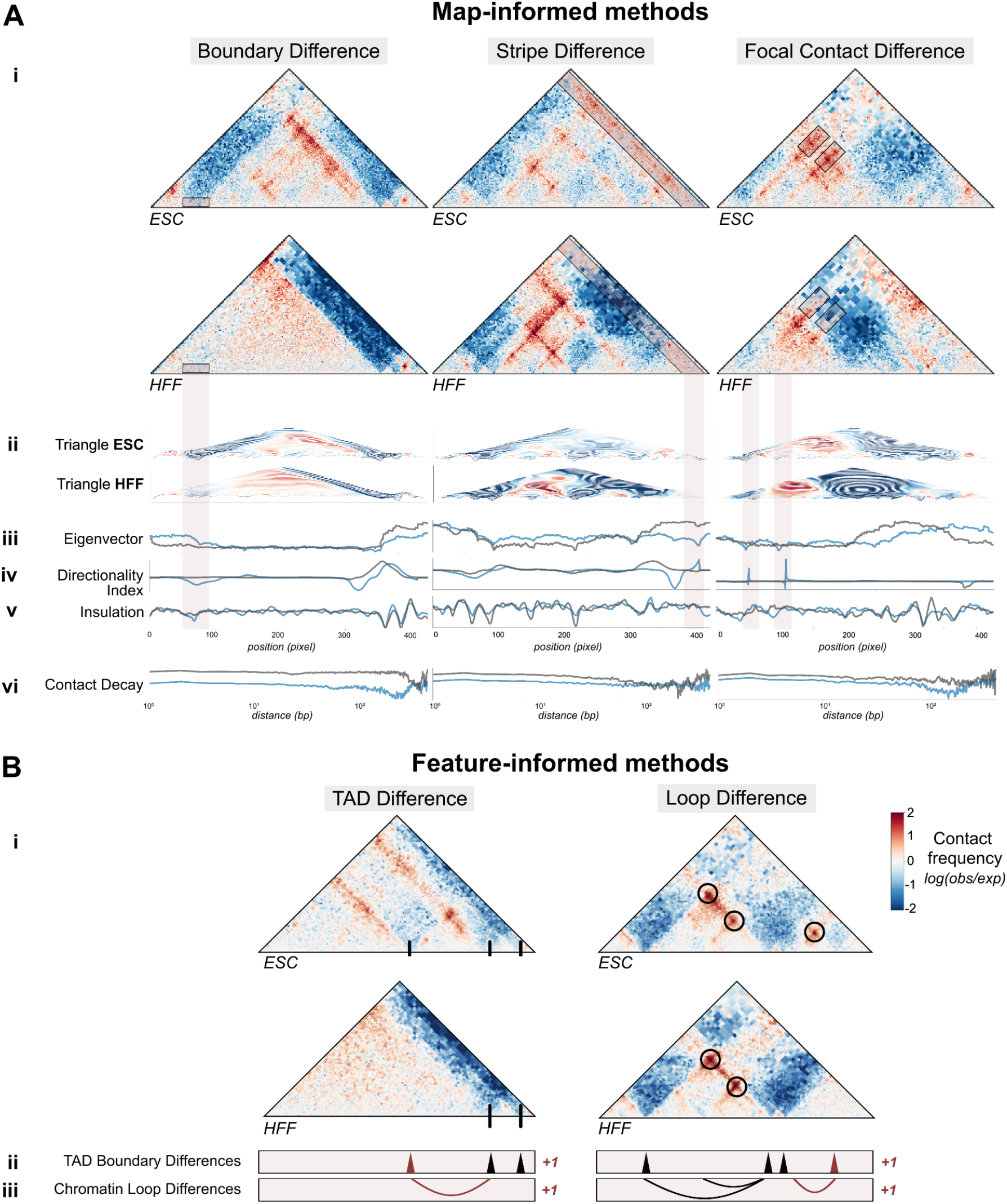
Map-informed and feature-informed methods capture differences in TAD boundaries, stripes, and loops. **A. *i.*** Examples of regions where contact frequency maps differ between HFF and ESCs across three structural changes: a lost TAD boundary (left panel), a lost stripe (middle panel), and lost loops (right panel), as marked by red boxes. ***ii-vi***. Tracks corresponding to each map-informed disruption score method are shown below for ESCs (blue) and HFF (gray). Tracks for methods in (ii. - v.) correspond to the coordinates of the contact maps, while contact decay in (v.) is plotted across genomic distance. **B. *i.*** Two loci in HFF and ESC with a boundary and loop change (GRCh38 `chr3:137129984-138178560` and GRCh38 `chr3:138702848-139751424`, respectively). ***ii.*** Applying a TAD boundary caller identifies a boundary change between cell types ***iii*.** Comparing chromatin loops identifies a genomic region with differential looping.

### Feature-informed methods prioritize changes to interacting chromatin regions

To evaluate comparison approaches based on TAD and loop calling methods, we chose two regions with differential structure between ESC and HFF maps (**Fig. 3B i**) and tuned the parameters of the cooltools TAD caller ^31, 32^ and the HiCCUPS loop caller ^28^ (**Supplemental Fig. 2**) ^35, 36^. As expected, the TAD caller correctly identified all three TAD boundaries visible in ESCs, including one that is lacking in HFF (**Fig. 3B ii**). Similarly, the loop caller identified a loop that is unique to ESCs (**Fig. 3B iii**). While these feature-informed approaches are biologically interpretable, they tend to be slower, address only one element at a time, and require additional parameter selection (**Table 1, Supplemental Table 1**). These methods also require a significance cutoff for initial feature calls, which may result in missed features of low signal. Additionally, most maps contain fewer than ten called features in a 1-Mb window, creating a small range of possible scores. Therefore, caution should be exercised when using these scores at a large scale, especially in maps without strong TADs or loops, where they can produce artificial results.

**Table 1:** Strengths, weaknesses, and suggested applications of disruption score methods. Trends and patterns across disruption scores summarized from statistical comparisons (**Fig. 4**), simulations (**Fig. 5**), and manual parsing of the most highly disruptive perturbations for each method (**Supp. Fig. 5**). While this summary is not exhaustive of all possible outcomes, it provides qualitative guidelines for users to make informed decisions when selecting a comparison method based on the scale and application of their research. We use green checks to indicate advantages and red X’s to indicate disadvantages for each method category: basic methods (blue), map-informed methods (orange), and feature-informed methods (green). Double signs represent strong patterns, while no sign indicates no pattern, and NA denotes that the method was not tested.

| Comparison method | Technical sensitivities |  |  |  | Application-based features |  |  |  | Specs |  |
| --- | --- | --- | --- | --- | --- | --- | --- | --- | --- | --- |
|  | Resistant to noise | Resistant to decreased resolution | Biased towards contrast maps |  | Sensitive to small structural changes | Sensitive to signal change at boundaries | Sensitive to loss or gain of boundaries | Sensitive to changes in TAD substructures | Needs tuning | Time & memory |
|  |  |  | High | Low |  |  |  |  |  |  |
| MSE |  |  | ✗ |  |  | ✓ | ✓✓ | ✓ |  | ✓ |
| Corr |  | ✓ |  | ✗ |  | ✓ | ✓✓ |  |  | ✓ |
| SCC |  |  |  | ✗ |  | ✓ | ✓✓ |  |  | ✓ |
| SSIM |  |  |  |  | ✓ | ✓ | ✓✓ | ✓ |  | ✓ |
| Eigenvector (corr)* | ✓✓ | ✓✓ |  |  | ✗ | ✓✓ | ✓ | ✗ | ✓ |  |
| DI (corr) | ✓✓ | ✓✓ |  |  | ✓ | ✓ | ✓✓ | ✓ | ✓ | ✓ |
| Insulation (mse) | ✓ |  | ✗ |  | ✓ | ✓✓ | ✓ | ✓ | ✓ |  |
| Insulation (corr) | ✓ |  |  | ✗ |  | ✓✓ | ✓ |  | ✓ |  |
| Contact decay (corr)** | ✓ | ✓✓ |  | ✗ |  |  | ✓✓ | ✗ |  |  |
| Triangle (mse)* | ✓ | ✓ | ✗ |  | ✓ | ✓✓ | ✓✓ | ✓ |  | ✗✗ |
| Triangle (corr)* | ✓✓ | ✓✓ |  | ✗ |  | ✓ | ✓✓ |  |  | ✗✗ |
| Loop calling | NA | NA |  |  | ✗ | ✓ | ✗ | ✓ | ✓ |  |
| TAD calling | NA | NA |  | ✗ | ✗ | ✗ | ✓ | ✓ | ✓ |  |

### *In silico* perturbation enables evaluation of contact map comparison methods at scale

Although differences between cell-types exist, 3D genome organization is often highly conserved^7, 30, 37^. To evaluate the performance of map comparison methods across a wider variety of possible changes in chromatin structure, we used an *in silico* approach to generate pairs of 1-Mb maps across the genome with a variety of perturbations. We applied Akita ^20^, a convolutional neural network that predicts genome folding from sequence alone, to generate contact frequency maps from sequences with and without a genetic perturbation likely to disrupt genome folding (**Fig. 4A**). We designed three types of perturbations: CTCF canonical motif insertions ^38^, endogenous CTCF motif deletions, and random 100 base pair deletions (**Methods**). In total, we produced 22,500 unique contact frequency map pairs on which to test all three types of methods. To enable large-scale evaluation, we applied the 11 methods and transformed their scores such that higher values indicate greater disruption of 3D organization and smaller values indicate more similar organization (**Methods, Supplemental Fig. 3**).

**Figure 4.**
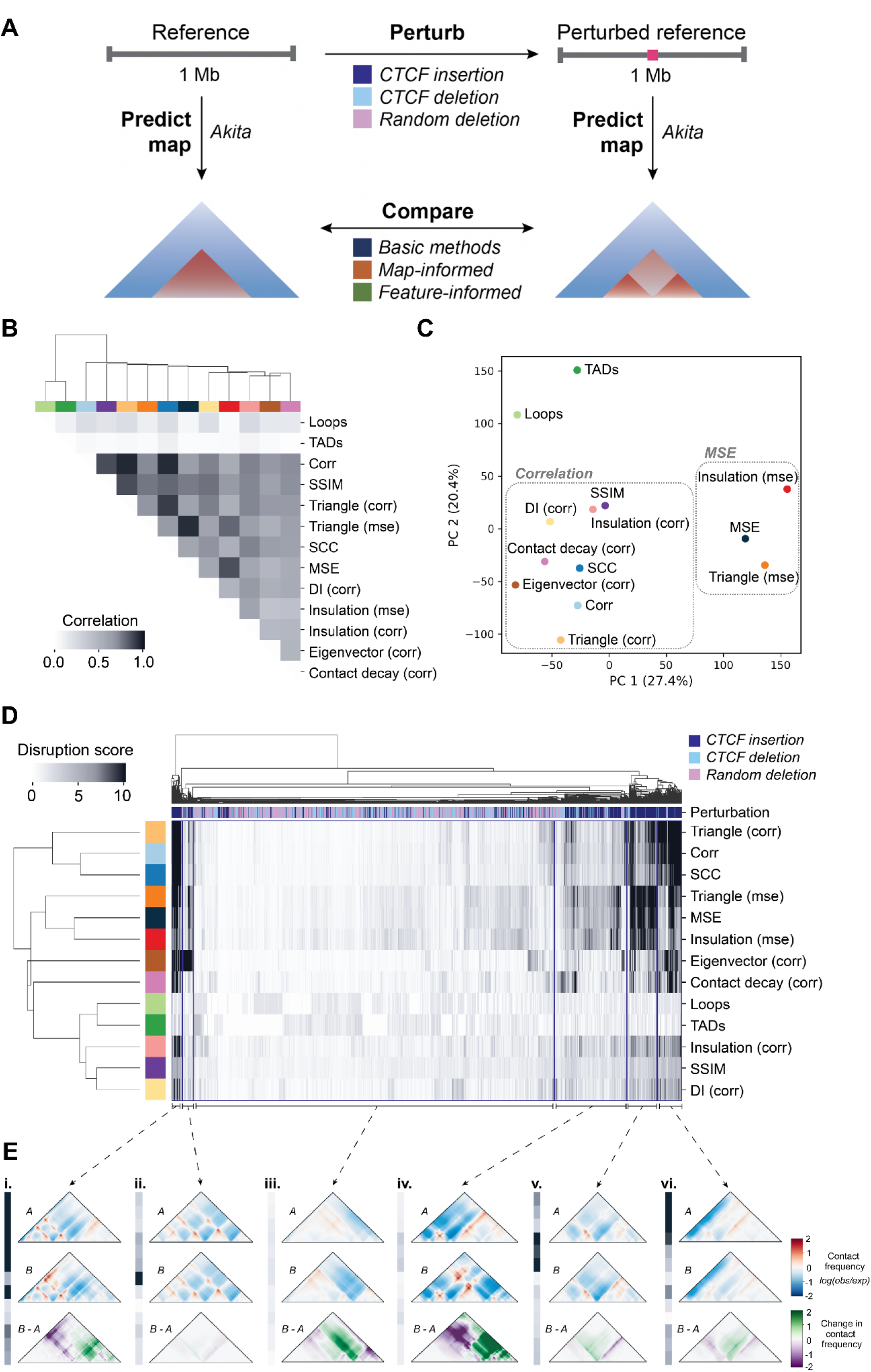
Comparison of disruption score methods. (**A**) Schematic describing the strategy for comparing *in silico* perturbed contact maps. Random ∼1 Mb windows of the human genome (GRCh38) are selected and input into Akita to predict chromatin contacts (left). The same window is also perturbed with a CTCF motif insertion, deletion, or random 100 base pair deletion. The resulting sequence is also input into Akita to predict chromatin contacts of this perturbed reference sequence (right). The perturbed and unperturbed maps were compared by applying the 11 basic, map-informed, and feature-informed methods. (**B**) Correlation matrix of the methods tested, where cells are shaded according to how well their scores correlated across perturbations. Concordance of the top-scoring perturbations (**Supp. Fig. 6**) also shows agreement between corr and SSIM, while highlighting that loops and triangle (corr) are quite concordant with other methods when considering only the top scores. Colors across the top of the heatmap identify the individual methods. (**C**) Principal component analysis of disruption scores of each method from perturbed map pairs. (**D**) Heatmap of normalized disruption scores across all methods and perturbations. The colored key along the top of the heatmap indicates whether the perturbation was a random deletion (pink), a CTCF insertion (navy), or a CTCF deletion (light blue). Method colors are the same as in (C). Four broad trends in disruption score patterns across methods are marked with brackets. (**E**) Representative example map pairs chosen from the groups identified in D: i. high scores across 5 methods; ii. low across all methods except for eigenvector (corr); iii. low scores across all methods; iv. low scores across methods but higher for MSE-based scores; v. high scores only for MSE-based scores; vi. high scores for correlation-based scores: triangle (corr), corr, and SCC.

We quantified the similarities and differences between methods by comparing the scores for all 22,500 *in silico* perturbations across all possible pairs of methods. We found that TAD- and loop-based scores are most different from the rest, as they only detect a specific type of change (**Fig. 4B**). Correlation-based measures (i.e., Spearman’s correlation, SSIM, and correlation of contact decay) cluster together distinct from MSE-based methods (i.e., MSE, triangle (MSE), insulation (MSE)). This result aligns with our initial observation that Spearman’s correlation and MSE often do not agree, especially across their top-scoring variants (**Fig. 2**, **Supplemental Fig. 4, Supplemental Fig. 5**). Principal component analysis (PCA) on the disruption scores shows similar clustering (**Fig. 4C**).

We next simultaneously clustered the perturbed map pairs and scores across methods to identify groups of perturbations that differentiate them (**Fig. 4D**). While all correlation-based methods exhibit similar behavior, insulation (corr), SSIM, and DI (corr) produce scores which are more uniformly distributed and less extreme across perturbations, highlighting the necessity of appropriate normalization when comparing across methods (**Fig. 4E i and vi**). We also find that perturbations created by CTCF insertion group together, as they are often the most disruptive of 3D organization. However, we observed substantial sub-structure within the cluster, reflecting differences in the behavior of scores on these maps. For example, cluster ***i*** is highly scored by all methods, and a representative perturbation example shows a variety of changes: gained loops, lost stripes, and boundary changes. The magnitude of changes in this set likely contributes to the universally high scores. Clusters ***iv*** and ***v*** are primarily composed of CTCF insertions, where scores are similar across most methods, but higher only for MSE-based methods. Profile ***v*** is the most dissimilar. Here, the representative map pair has minimal structural differences but extreme contrast, suggesting that this cluster is defined by examples of high dynamic range that are over-prioritized by MSE-based methods (**Fig. 4E v**).

We further compared methods by quantifying how well the top-ranked maps agree across methods. Some methods have high overlap (**Supplemental Fig. 5**, **Supplemental Fig. 6**). For example, 85% of map pairs are ranked in the top 5th percentile for both SCC and Spearman’s correlation, indicating some general agreement in the methods. However, many methods have minimal overlap, suggesting they prioritize different features. For example, only 32% of the top 5th percentile of maps ranked by insulation (MSE) and SSIM are shared. Finally, we applied methods to map pairs selected to represent a range of effect sizes and confirmed all methods are sensitive to large changes and insensitive to small changes (**Supplemental Fig. 7**).

**Figure 5.**
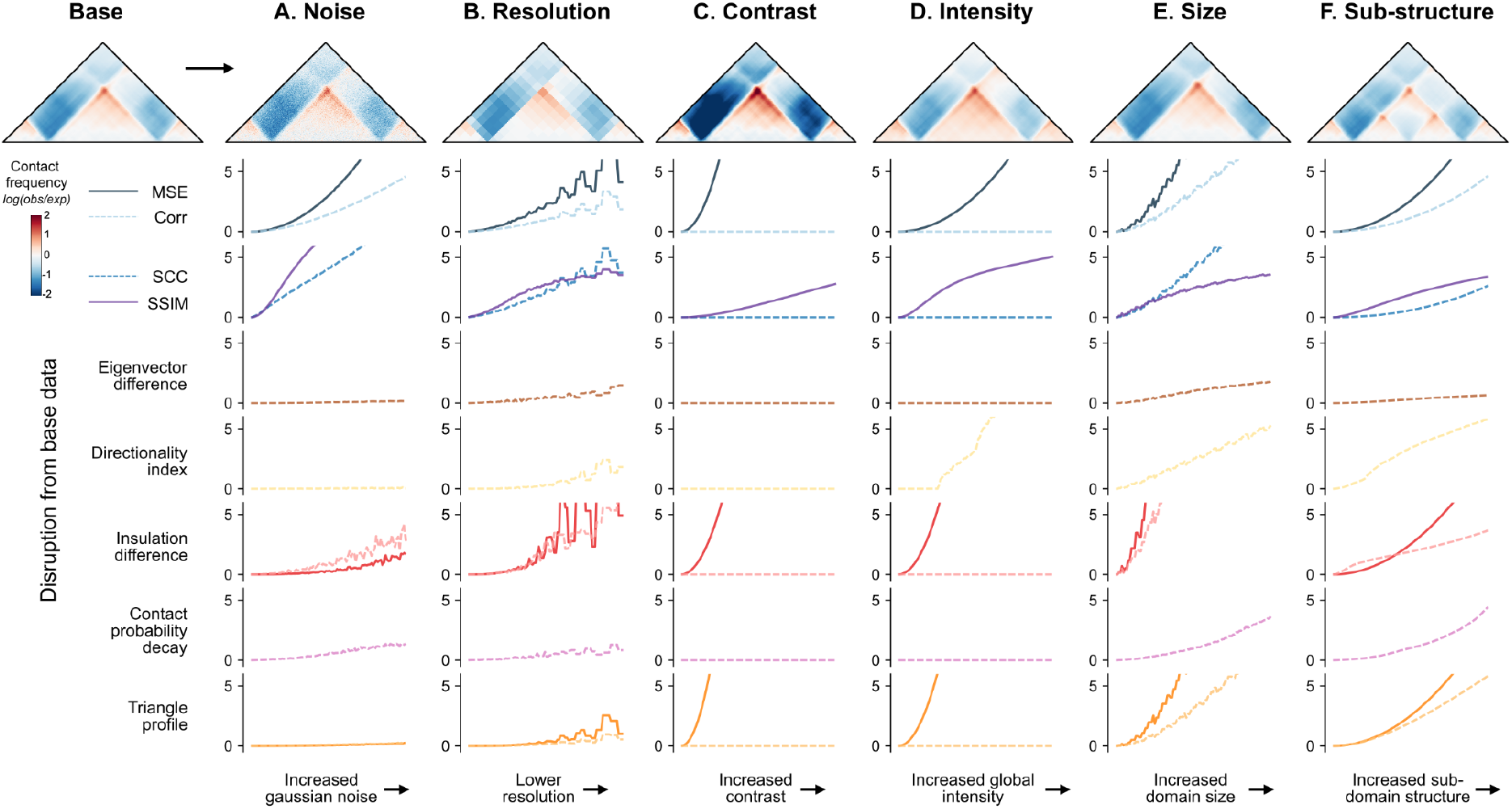
Simulated contact frequency maps with controlled perturbations estimate disruption score method sensitivities. Normalized disruption scores are plotted for a simulated contact frequency map containing a TAD across 6 types of perturbations, plotted on the x-axis. Each perturbation was added at 100 different degrees. The images shown correspond to the final degree–the maximum perturbation added. Line plots show disruption scores from comparing the original map (top left corner) to each perturbed map. Maps corresponding to the incremental increases in perturbation are shown alongside the changed scores in **Supp. Fig. 8**. (**A**) Noise is added by introducing random values drawn from a Gaussian distribution to the maps; (**B**) Resolution is lowered by increasing bin size; (**C**) Contrast is applied by increasing the range of the signal; (**D**) Intensity is increased globally by adding a constant to all values; (**E**) Size is increased by slightly enlarging the domain width; (**E**) A sub-structure is added by gradually incorporating a new boundary at the center of the existing TAD.

### Simulation studies quantify method sensitivity

Our *in silico* screen produced a diversity of structural alterations, often affecting multiple aspects of the map. For instance, a CTCF site insertion can both create a new TAD boundary and alter overall contact intensity. To disentangle how each method responds to changes in particular map features, we generated simulated maps and synthetically altered a single variable at a time. We then measured the sensitivity of each score to each specific change. As a template, we created a contact frequency map with two CTCF motifs forming a TAD and used this canvas to simulate both biologically meaningful changes (e.g., change in TAD size, substructure, or intensity) and technical artifacts (e.g., change in noise or resolution) (**Methods**). For each change, we gradually increased the strength of the perturbation across 100 maps and subsequently applied scoring methods (**Supplemental Fig. 8**).

Each method responded differently across the simulated changes (**Fig. 5**). Steeper curves represent high sensitivity to the perturbation, while flatter curves represent less sensitivity. We find that basic methods are most sensitive to technical variations, such as increased noise and decreased resolution, while map-informed methods are most robust (**Fig. 5A-B**). As expected, correlation-based methods are unaffected by changes in contrast and intensity, while MSE-based methods are highly sensitive **(Fig. 5C-D**). All methods except eigenvector difference identify TAD size and sub-structure changes. However, some prioritize certain types of organizational changes over others (**Fig. 5E-F**). For example, insulation difference and triangle profile are sensitive to boundary changes, while directionality index highlights new boundaries but is less effective in identifying changes to existing boundaries. We synthesized these results along with findings from *in silico* perturbations in the **Guidelines** to provide recommendations based on the intended application.

### Guidelines

Our study assessed the effectiveness of 11 existing and new methods for comparing 3D genome contact maps (**Supplementary Table 1**). Although there were similarities between the top-scoring variants of most methods, our results indicate that they differ substantially in their sensitivity to biological and technical variation (**Supplemental Fig. 6**). We summarize these findings and guidelines in **Table 1**.

All of the methods can identify structural changes, such as changes to domain size or the addition of substructure, but to varying degrees. Of the basic methods, MSE and SCC more readily identify subtle organizational changes. Among the map-informed methods, insulation difference and triangle difference are the most effective at identifying changes in both existing and new domain boundaries. Directionality index highlights new boundaries or substructures but less readily identifies changes in existing boundaries. Eigenvector difference and contact probability decay are the least sensitive to small-scale organizational changes, but prioritize larger-scale changes in the overall structure of the map. These statistics have been deployed primarily for identifying differences at the scale of compartments and whole chromosomes, so it is not surprising that they are not sensitive to map differences within 1-Mb windows.

In general, the new map-informed methods we proposed are concordant with basic methods and each other, especially when comparing the top 5% of scores genome-wide (30%-80% of examples are shared; **Supplemental Fig. 6**). Triangle difference stands out among our newly implemented methods as highly concordant with other methods and able to detect a variety of map differences, but it is also the slowest (**Supplementary Table 1**). Insulation difference is faster and also fairly concordant with other methods. Most top 5% map pairs called by other methods are also high-scoring with loop calling, but not TAD calling. Loop calling also identifies many additional map pairs that are not in the top 5% of other methods.

Correlation-based methods are insensitive to changes in contrast and intensity, while MSE-based methods are highly sensitive to these changes. In contrast, map-informed methods summarize maps across a feature track and are therefore more robust to these changes. One notable exception is insulation difference, which is more sensitive to resolution changes that obscure domain boundaries. Some map changes, such as contrast or intensity, may either be biologically meaningful or a consequence of technical variability, depending on the scenario. The basic methods, especially MSE, SCC, and SSIM, are particularly sensitive to technical variation such as increased noise and decreased resolution. SSIM falls in between. We also note that MSE is by far the fastest approach (**Supplementary Table 1)**. All others require less than 10 seconds per thousand calls, aside from eigenvector difference and triangle track, which can be accelerated by decreasing the resolution of the maps prior to comparison.

We recommend using multiple methods in tandem. We find that there is no “one size fits all” metric that best identifies every feature of interest in a chromatin contact map. Researchers should consider the intended application and the types of changes that are meaningful when selecting the most effective and relevant metrics. We recommend first applying basic methods as an initial screen to identify the most disrupted maps, especially when evaluating large datasets. Using both correlation- and MSE-based scores will help mitigate biases of each. We next suggest applying a map-informed method, such as triangle or insulation difference, to a subset of disruptive perturbations to gain insight into the types of changes present. Finally, feature-informed methods can be used to explore TAD and loop gains/losses and to develop mechanistic hypotheses.

### Code

Our codebase is publicly available to enable researchers to easily test and apply all 11 approaches to their own research questions. The code is written in Python and is accompanied by documentation and tutorials to help users get started. The methods have flexible hyperparameters and can be run simultaneously on one dataset, making it easier to compare the results of different approaches and select the most appropriate methods. To aid in interpretation of the methods, we also provide guidance on how to visualize map-informed and feature-informed approaches across contact matrices. Overall, our codebase provides a valuable resource for researchers who wish to apply multiple methods to their own datasets and rank pairs of maps based on their differences.

## Discussion

In this study, we evaluated and compared the behavior of 11 methods for quantifying differences between pairs of 3D contact maps, including many methods that have not been previously used for this application. We introduced insulation difference, eigenvector difference, and contact decay difference, as well as the new triangle comparison method, which is robust to noise while capturing structural differences between maps. We found that the choice of scoring function can have a significant impact on the conclusions drawn from the data, and therefore suggest that multiple comparison metrics should be used when seeking biological insights into the function of the 3D genome.

Several limitations should be considered when evaluating our results. While we consider a range of experimental, predicted, and simulated maps, our findings may not apply to other experimental conditions, such as single-cell contact matrices or other scenarios in which maps have a high level of noise and/or sparsity. Additionally, some of the methods we evaluated have variables that can be tuned to optimize performance in a given context (**Supplementary Text, Supplementary Table 1**). We only tested one TAD caller and one loop caller to examine their general utility ^35, 36^. Finally, we did not directly address the problem of identifying a threshold beyond which the differences should be considered biologically or statistically significant. One could apply previously proposed ^6, 25, 39^ and novel thresholding methods to the ranks computed with scoring methods to define a significant set of map pairs.

Our work provides useful guidelines for scoring contact maps that will enable further discovery into the mechanisms of the 3D genome. We provide a codebase of methods for flexible and fast scoring across contact maps under a unified framework. The experiments we performed as a part of this study, such as the *in silico* deletion and insertion of thousands of CTCF motifs genome-wide, provide a useful dataset for evaluating diverse biological questions or utility as controls for the level of 3D genome variation expected based on CTCF and random perturbations. We anticipate that incorporating methods with stronger biological interpretability, like those evaluated here, may further improve machine learning methods for predicting contact maps. Overall, by developing novel and more robust scoring functions, our study provides a foundation for analyzing contact maps at scale.

## Code and Data Availability

All original code and resulting data from *in silico* screens have been deposited at –https://github.com/pollardlab/contact_map_scoring.

## Acknowledgements

We gratefully acknowledge members of the Pollard lab for project feedback, as well as Vijay Ramani, for proposing eigenvector difference. We additionally thank Mike Keiser for feedback and support.

## Methods

### Datasets

#### Experimental maps

Maps of 3D chromatin contact are represented as 2D matrices of pairwise interaction frequencies. Regions of maps with high values indicate genomic loci with a high frequency of interaction in physical space, on average. Following experimental Hi-C, maps begin as raw read counts, which are subsequently balanced and normalized to reflect log(observed/expected) contact frequencies ^40^.

Experimental data considered in this study from HFF and ESCs were preprocessed as training datasets for the Akita model^11, 20^. Specifically, these high-quality Micro-C datasets were normalized with genome-wide iterative correction (ICE), adaptively coarse-grained, normalized for distance-dependent decrease in contact frequency, log clipped to (−2,2), linearly interpolated to fill missing bins, and convolved with a 2D Gaussian filter for smoothing. Processing maps ensures consistency across the experimental data and computational predictions since we do not evaluate raw experimental read counts.

#### Predicted maps

To effectively compare contact maps at scale, we generated a dataset of thousands of maps predicted from *in silico* CTCF motive insertions, CTCF motif deletions, random 100 bp sequence insertions, and random 100 bp sequence deletions. These alterations were passed into Akita^20^, a model predicting genome folding from sequence, to generate pairs of maps with structural rearrangements. We first curated sequences for insertion. CTCF motif sequences were randomly selected from annotated CTCF sites in the reference genome from the hg38 build of the JASPAR database^38^. Random 100 bp fragments were also selected from chromosome 1 for insertion. Both the CTCF and random sequences were inserted into the center of 1-Mb of DNA with start locations randomly selected from chromosome 1. Akita requires a fixed input of 2^20^ bp. Additional sequence was trimmed from the 3’ end, such that the final sequence remained 1-Mb. To curate deletions, we again selected random CTCF sites from JASPAR, pulled the surrounding 1-Mb of DNA, removed the motif sequence, and pulled in additional sequence from the 3’ end such that the entire sequence remained 1-Mb in length. The same strategy was applied to randomly selected 100 bp fragments for deletion. All generated 1-Mb genomic query sequences were filtered to exclude overlap with ENCODE blacklisted regions^41^. For each perturbation, both the original genomic sequence and the perturbed sequence were provided to Akita, resulting in two predicted 448x448 contact maps where the resolution of each pixel is 2048 bp representing a total length of ∼1 Mb (2^20^) of DNA sequence ^20^. This dataset consists of 7,500 matched contact maps for each category of perturbation for 30,000 total map pairs. Random 100 bp insertions were generally excluded from analysis, as they had almost no effect.

#### Simulated maps

To generate simulated maps, we initially generated predicted maps with Akita from random DNA sequence. Predicted maps still showed minimal structure from randomly occurring CTCF-like motifs. Sequence matches to the forward and reverse canonical CTCF motif^38^ were therefore shuffled to produce a predicted blank canvas map devoid of all higher-order folding patterns. Structure was reintroduced to simulated maps by inserting forward and reverse CTCF motifs ¼ and ¾ through the random DNA sequence, producing TAD-like boundaries. We tuned simulated parameters as described below.

● *Noise:* Gaussian noise was added to the maps with a standard deviation ranging from 0 (no added noise) to 0.2.
● *Resolution:* The original 448x448 map was downsampled ranging from a resolution of 2,048 bp (original resolution) to 50,972 bp.
● *Contrast:* Pixel intensities of the contact map were multiplied by a scalar ranging from 1 (no increase in contrast) to 2.
● *Intensity:* A scalar value ranging from 0 (no addition) to 0.2 was added to all pixels in the contact map.
● *Size:* The size of the substructure within the map was increased by resizing the original map by a scalar and trimming the matrix back down to the original dimensions. Map sizes were increased by a factor of 1 (no resize) to 1.1.
● *Substructure:* An additional map was created by introducing CTCF halfway into the random sequence to produce an additional boundary. The original map was combined with the substructure map with a multiplier ranging from 0 (no added structure) to 1 (total added structure).

Visualizations of these changes can be found in **Supplemental Fig. 8.**

### Benchmarking methods

#### Adapting new methods

Triangle profile is a novel scoring method. Directionality index^30^, PCA^29^, insulation^31^, and contact decay^9^ are established methods for analysis on individual Hi-C maps, but have not previously been used to to score pairs of maps. For map-motivated and feature-motivated methods, it is possible to plot the scoring method results along the length of the map, or on the map itself, as seen in Figure 3. The common behavior across maps with a small change, a large change, and no change is illustrated in **Supplemental Fig. 7**.

#### Comparing contact maps

We applied all comparison metrics to pairs of experimental, predicted, and synthetic maps. For details regarding how each metric is computed, see **Supplemental Text**. Any missing values were masked prior to evaluation and not considered by the comparison metrics. Scoring method implementations can be found within scoring.py in the codebase. MSE, Spearman’s rank correlation coefficient, and Pearson correlation coefficient were applied to map-informed methods to collapse two 2D tracks into a scalar value. Pearson correlation behaved almost identically to Spearman’s rank correlation, and therefore was excluded from analysis (**Supplemental Fig. 1)**. For computationally intensive methods, we reduced the resolution of the input from 2kb to 10kb to speed evaluation time across thousands of comparisons.

To ensure that scores across approaches are comparable, we flip some methods such that higher values indicate greater disruption and smaller values indicate more similar maps. For methods like correlations, we use 1 - correlation such that a perfect correlation (1) is flipped to mean no difference (0). For all the results, we provide raw scores and normalized scores so that it is easier to interpret how a raw score for one method compares to a raw score of another method. We additionally scale all values by the mean score of all random 100 bp deletions using Akita, which we find to have minimal impact (**Supplemental Fig. 3**). For example, a raw MSE of 0.0065 and a raw 1 - pearson correlation of 0.036 both correspond to the same normalized score of 2. That is, a disruption of that magnitude corresponds to 2 times the average disruption of a 100 bp deletion.

For loop and TAD callers, we quantify the ratio of changed (e.g. added or lost) features (TADs or loops) to extend these approaches and generate a single score for each pair of maps.

#### Method parameters

The following methods required no adjustable input parameters: mean squared error, Spearman’s rank correlation coefficient, and pearson correlation coefficient, SSIM, SCC, contact decay, eigenvector, and triangle correlation. We describe tunable parameters choices for the remaining methods below. We did not optimize tunable parameter choices but instead selected default choices from existing approaches. Results from alternative parameter selection are demonstrated in **Supplemental Fig. 2** and **Supplemental Fig. 9**.

Insulation:

*window_size*=10: size of the diamond-shaped window considered
Directionality index:

*window_resolution*=10000: resolution of sliding window in bp
*replace_ends*: replaces ends of DI track with 0s
*buffer*=50: how far from the track ends to replace with 0
Loop difference:

*p*=2: the width of the interaction region surrounding the peak
*width*=5: the size to get the donut filter
*ther*=1.1: the threshold for the ratio of center windows to the donut filter and lower left filter
*ther_H*=1.1: the threshold for the ratio of center windows to the horizontal filter
*ther_V*=1.1: the threshold for the ratio of center windows to the vertical filter
*radius*=5: the upper bound of distance of two loop points considered as same
TAD difference:

*window_size*=5: size of the diamond-shaped window
*ther*=0.2: the threshold for TAD boundaries
*radius*=5: the upper bound of distance of two TADs considered as same

## Supplementary Information

### Supplemental Figures

**Supplemental Figure 1.**
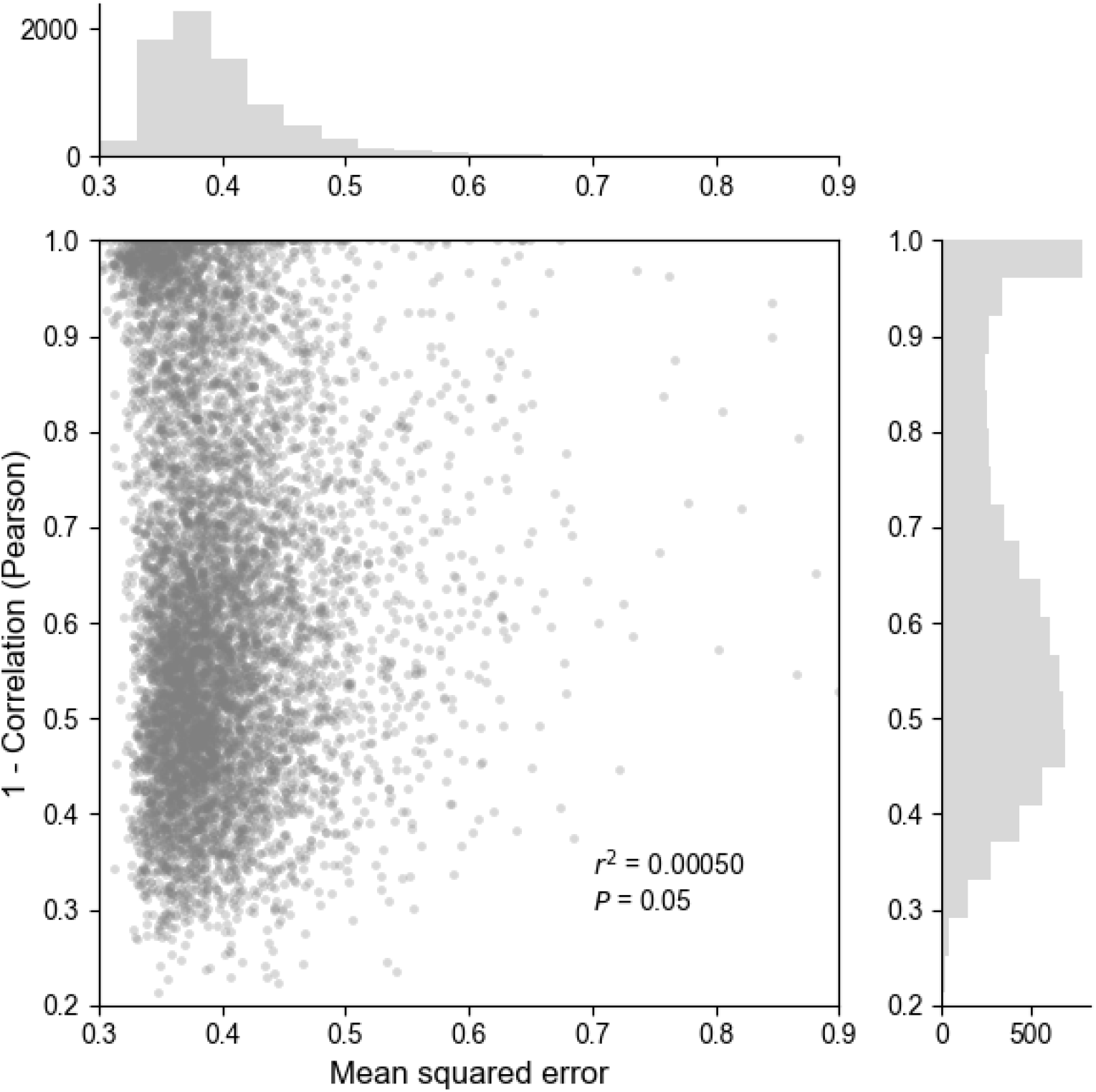
Pearson correlation versus mean squared error comparisons of contact maps. Mean squared error (MSE) and Pearson correlation coefficient calculated across the genome on experimental contact maps from embryonic stem cell (ESC) and human foreskin fibroblast (HFF). Each point represents a comparison between maps from HFF and ESC cell types (n = 7840 windows). There is a weak relationship between the Pearson correlation and MSE (r^2 = 0.0005, P = 0.05)

**Supplemental Figure 2.**
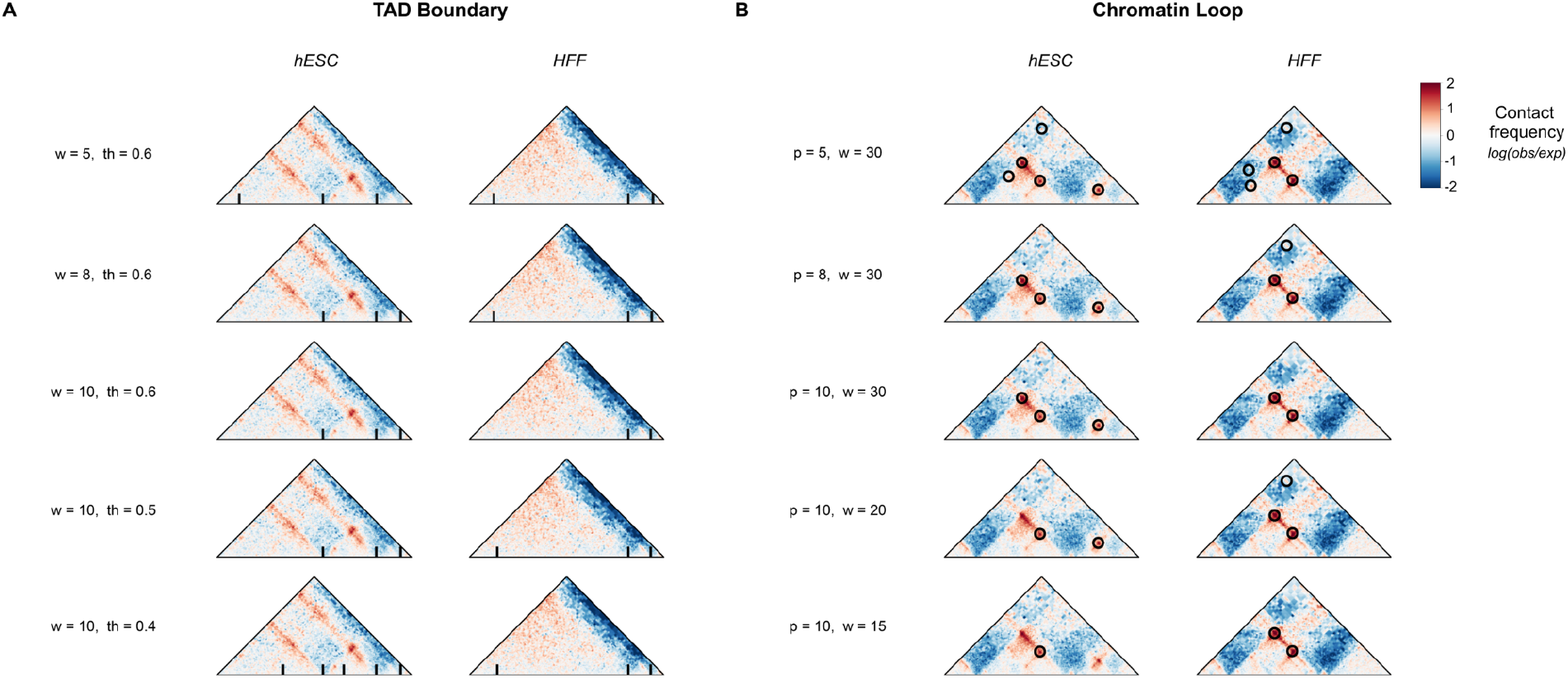
Sensitivity of TAD and loop caller on parameter shifts. (**A**) TAD boundaries (highlighted with black bar) called with different sizes of diamond-shaped window (*w*) and thresholds of insulation scores (*th*). (**B**) Chromatin loops (highlighted with black circle) identified using different sizes of center window (*p*) and donut filter (*w*). Example maps used here are the same as in **Fig. 3Bi**.

**Supplemental Figure 3.**
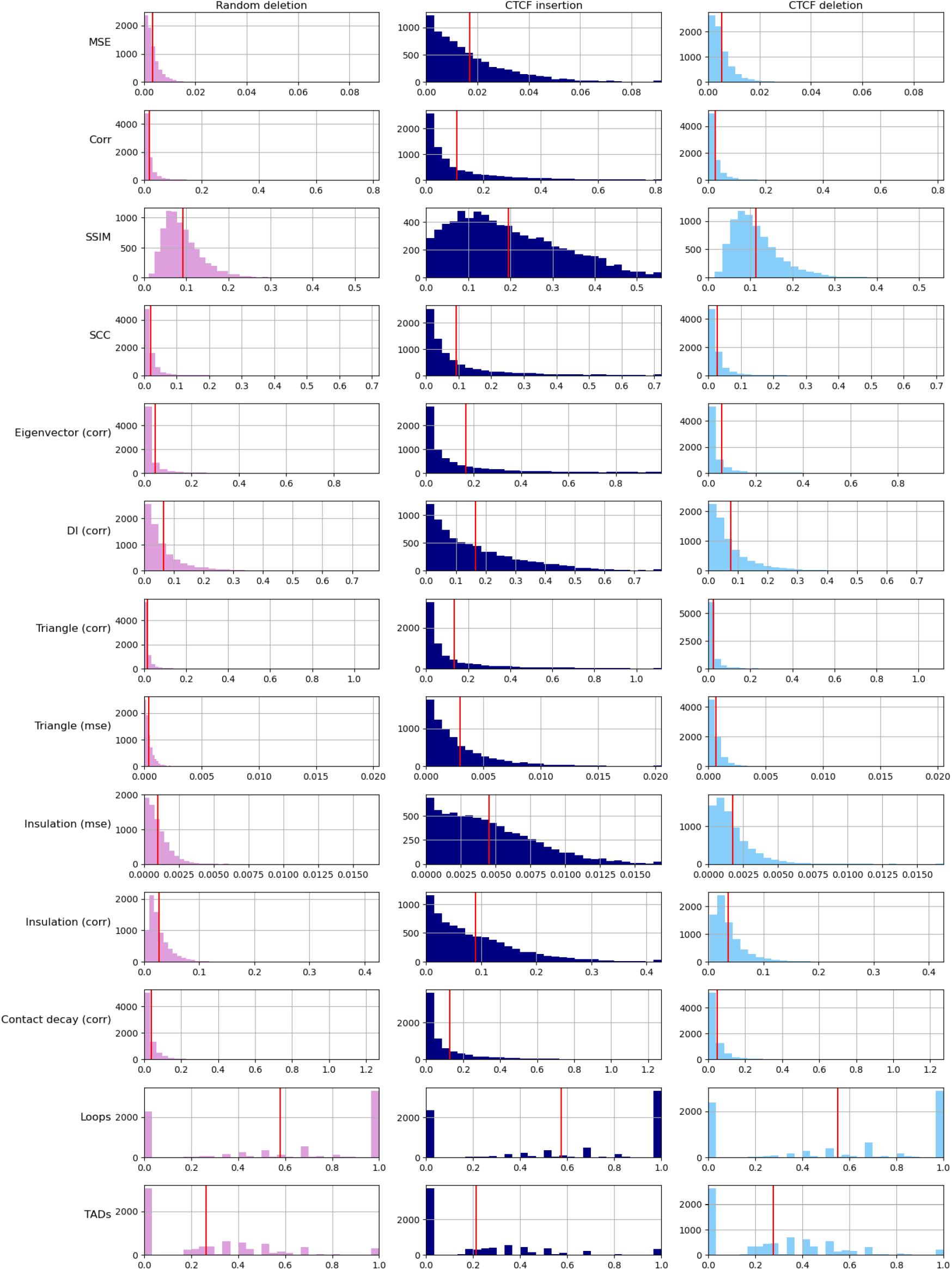
Score distributions of random deletions, CTCF deletions, and CTCF insertions. Each disruption score method (rows) produces a different range and mean (red line) across scores produced. Histograms show the raw scores comparing maps produced by 7500 random 100 bp deletions (left), 7500 CTCF insertions (middle), and 7500 CTCF deletions (right). To enable comparisons between the different scores, the main text figures report scores standardized to the mean disruption produced by a random 100 bp deletion. Thus, both an MSE-based disruption score and correlation-based disruption score describe that the maps are twice as different as the average 100 bp deletion.

**Supplemental Figure 4.**
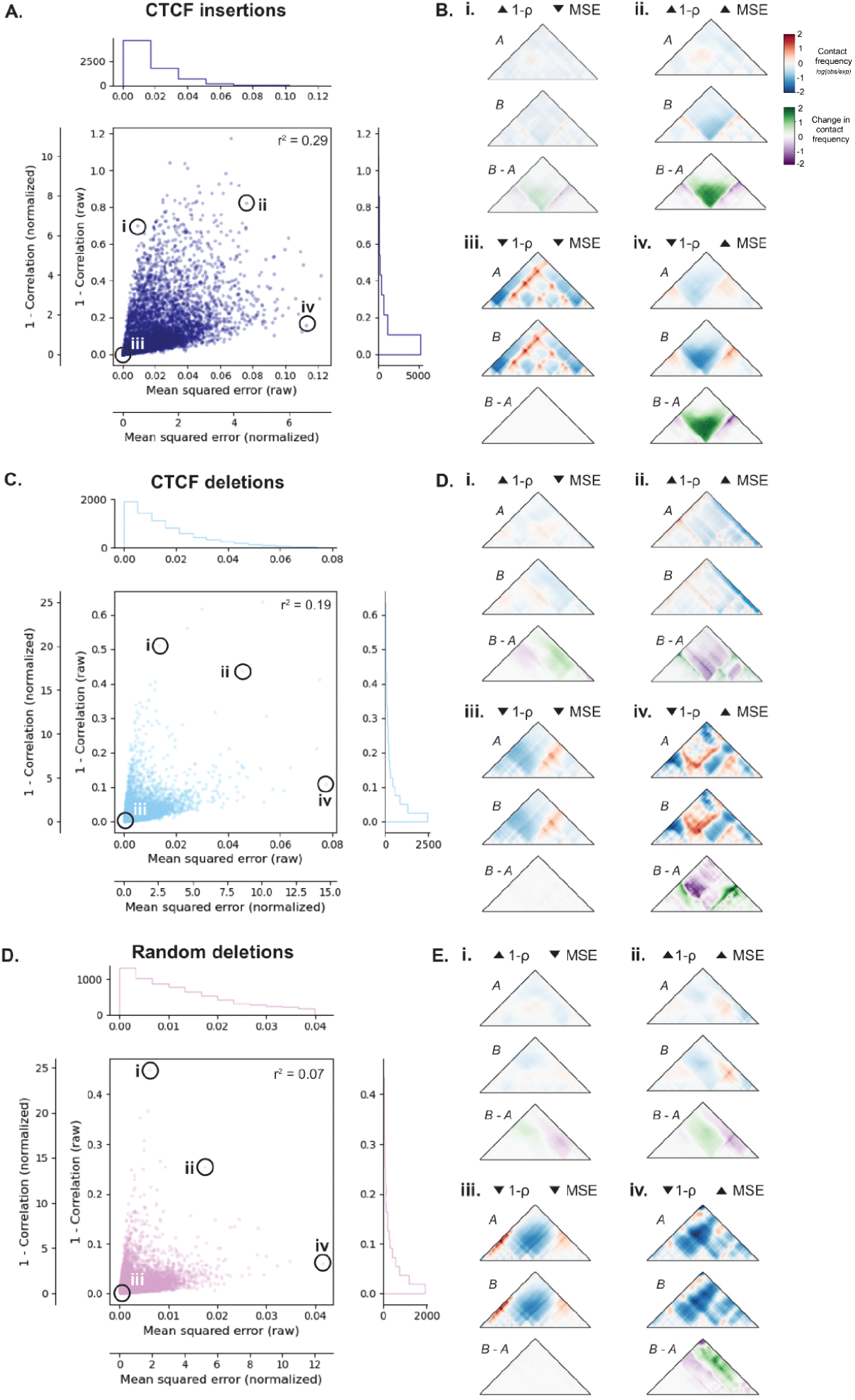
Basic methods to compare contact frequency maps rank maps differently on *in silico* perturbations. Mean squared error (MSE) vs Spearman’s correlation (ρ) scores across an *in silico* screen of 7,500 map pairs with and without CTCF insertions (A-B), CTCF deletions (C-D) and random 100 bp deletions (E-F), similar to **Fig. 2.** We plot 1 - ρ such that higher values for both methods reflect increasing differences between maps. MSE versus Spearman’s correlation are plotted where each point represents a comparison between a reference and perturbed map (**Fig. 4a**). Normalized scores are divided by the mean of the distribution of random deletions. Across all three perturbations, there is a weak relationship between the two disruption scores (r^2 = 0.29, 0.19, and 0.07 for CTCF insertions, CTCF deletions and random deletions, respectively). The relationship is strongest for CTCF insertions, for which scores are highest, followed by CTCF deletions, which have the next highest scores. Yet perturbations with at least one high score are not always concordantly scored. Examples of extreme scores for each perturbation are shown in B, D, and F in panels i through iv, illustrating that perturbations with high MSE and low 1 - ρ are consistently maps with high contrast, while low MSE and high 1 - ρ perturbations are maps with overall low contrast.

**Supplemental Figure 5.**
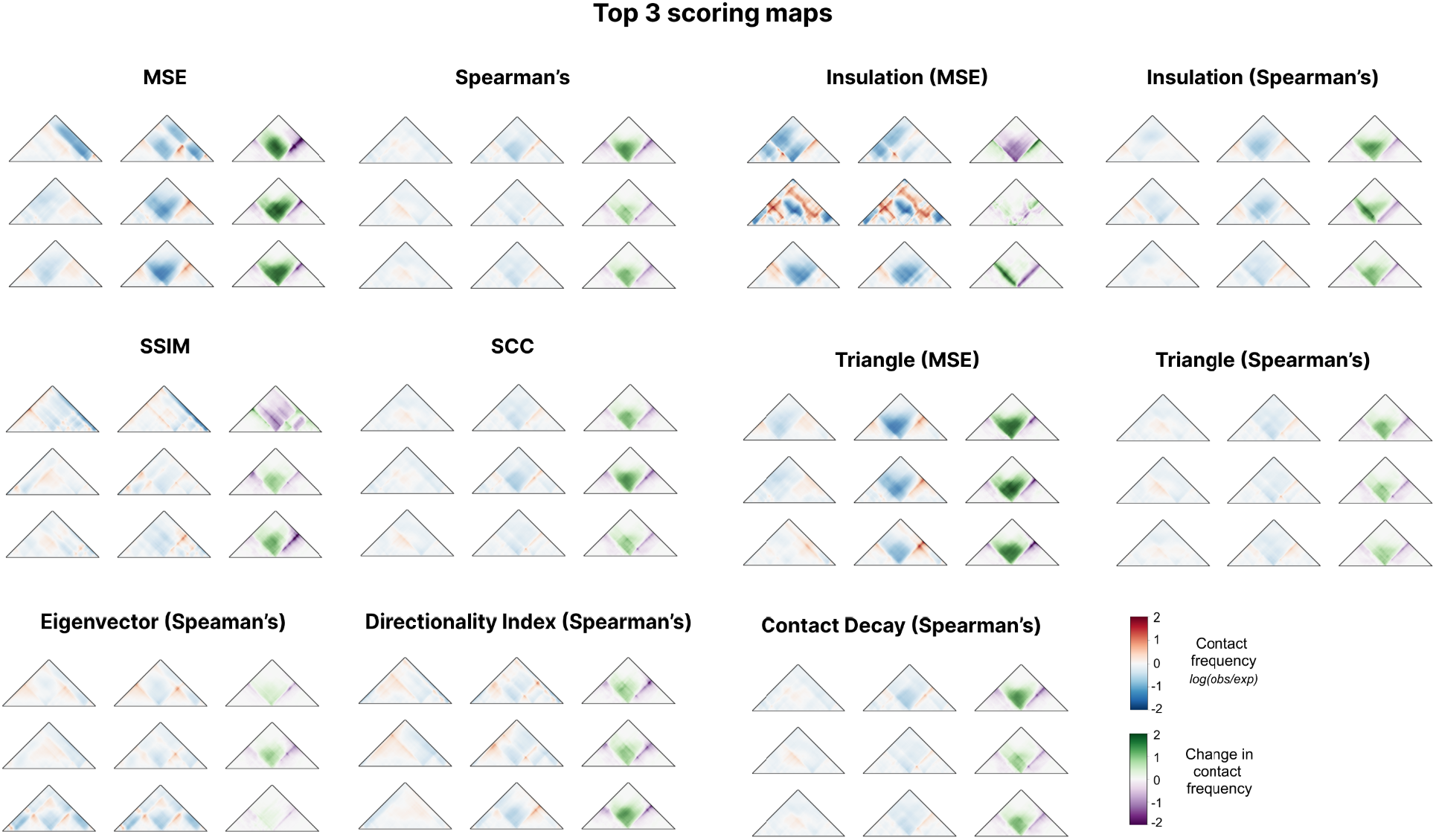
The three most disruptive map pairs of each scoring method. For each example row, the unperturbed map is shown on the left, the perturbed map is shown in the middle, and the difference between the two maps is shown on the right. The top three disruptive maps were chosen across the *in silico* screen.

**Supplemental Figure 6.**
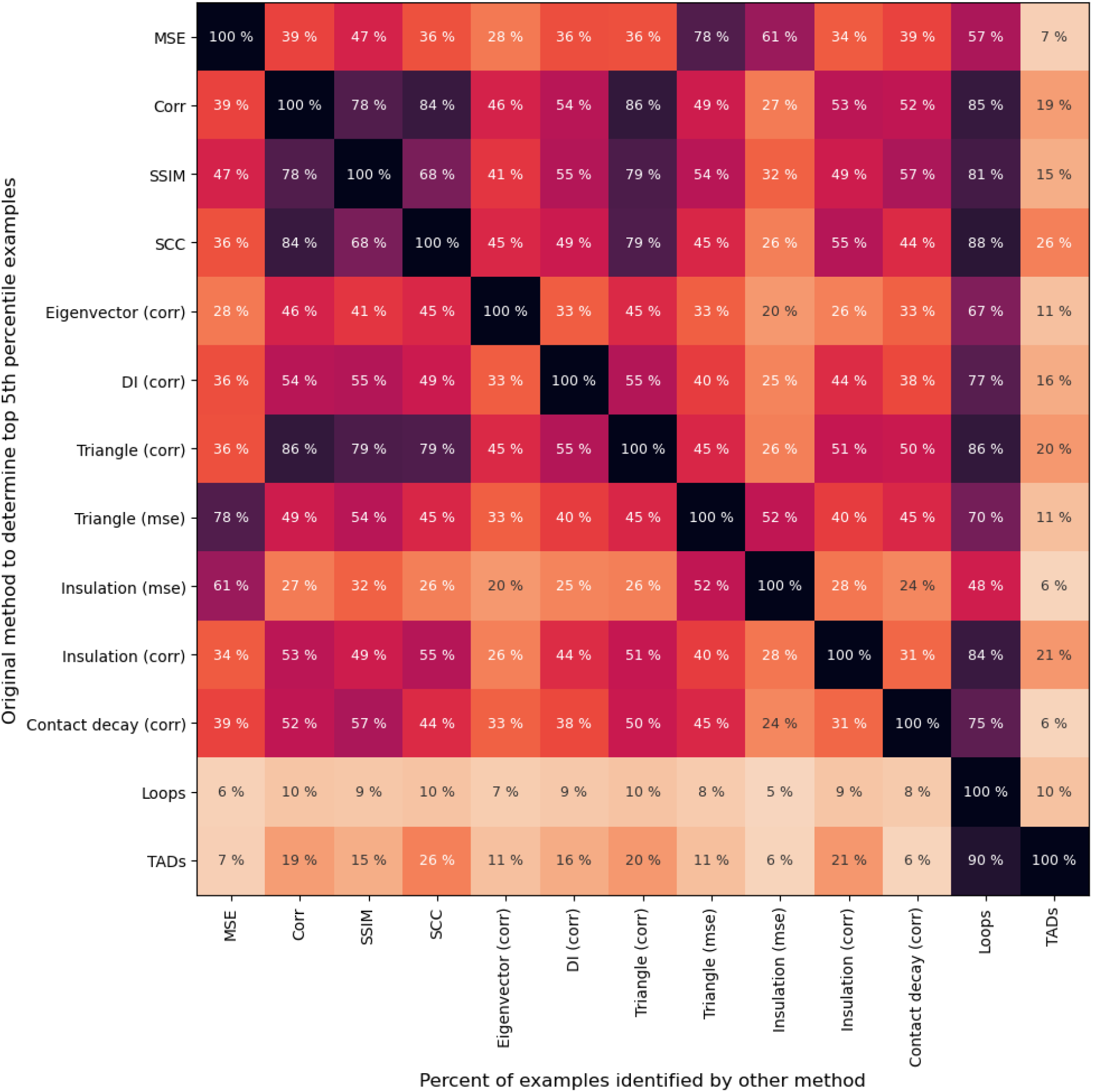
Overlap of the most disruptive map pairs identified by each scoring method. Each cell in the heatmap represents the percentage of map pairs that are above the 5% cutoff for the method in row and above the 5% cutoff for the method in the column. Darker colors indicate higher concordance for the top scoring loci. The heatmap is symmetric except for Loops and TADs. The imbalance of these two methods is caused by multiple map pairs that have scores equal to the 5th percentile, which results from methods producing low counts of discrete values.

**Supplemental Figure 7.**
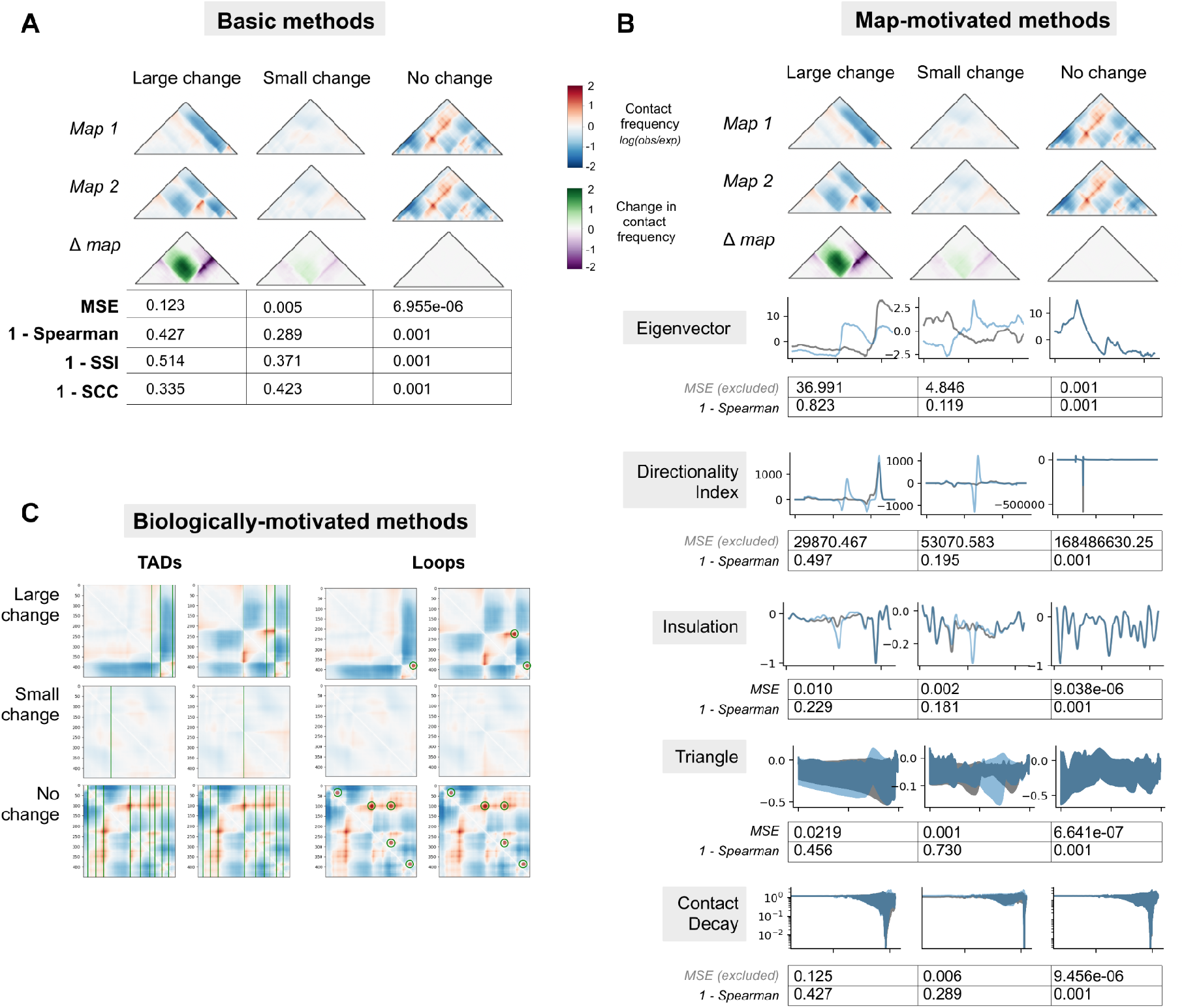
Scoring metrics on contact map pairs with large, small, and minimal changes. (**A**) Basic method scoring results across three example loci with a large, small, and minimal change upon CTCF motif insertion. (**B**) Map-motivated scoring results across three example loci. Raw tracks are shown for each measurement and the MSE and Spearman’s correlation between the tracks are shown below. (**C**) Feature-informed scoring examples across three example loci with a no change, a minimal change, and a large change to folding.

**Supplemental Figure 8.**
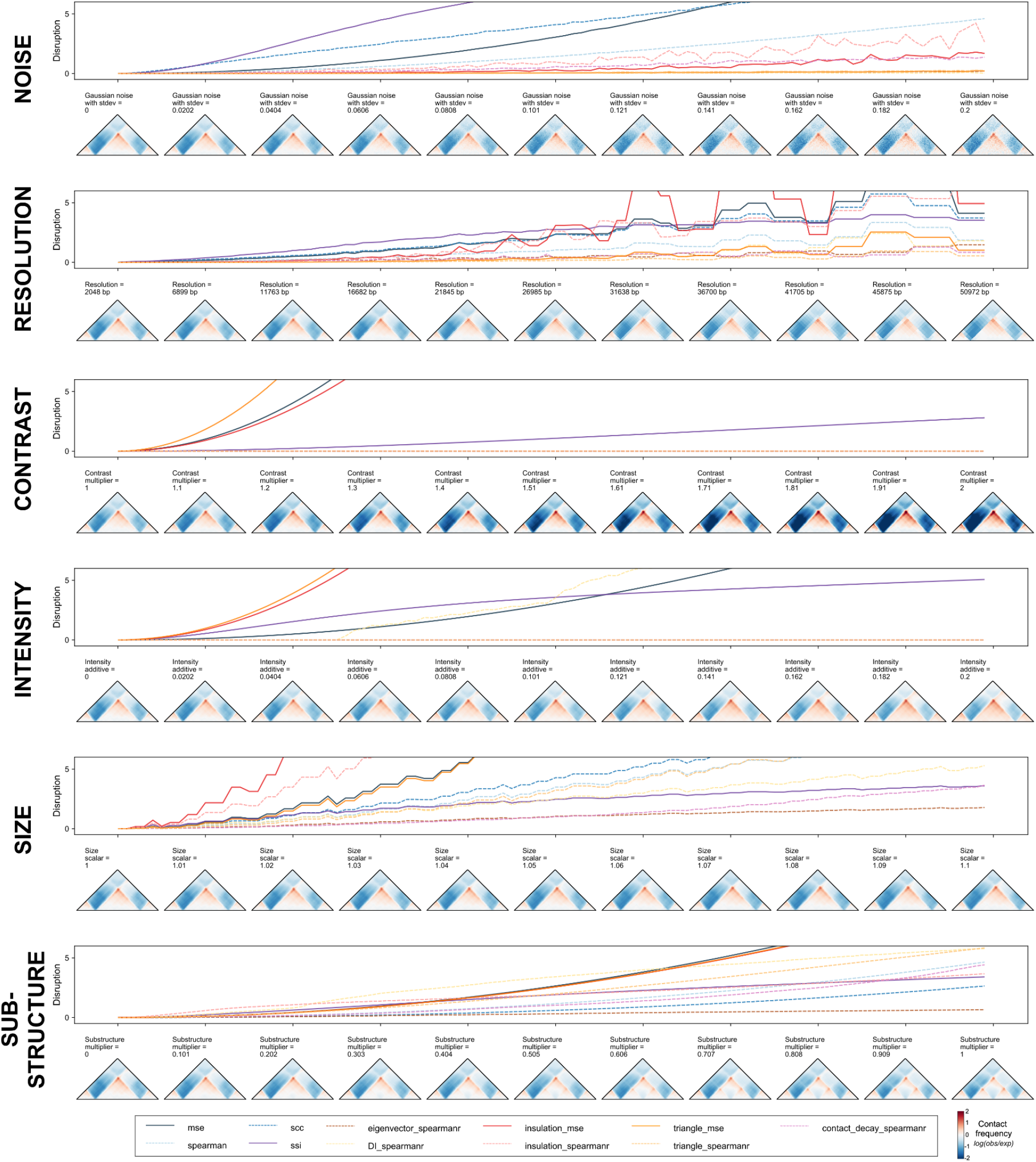
Changes of disruption scores with gradual increases in perturbations. Each subpanel shows the changes of disruption scores (top row) and contact maps (bottom row) against the incremental changes in a technical or biological variation. The colors of the scoring metric are the same as seen in **Fig. 5**.

**Supplemental Figure 9.**
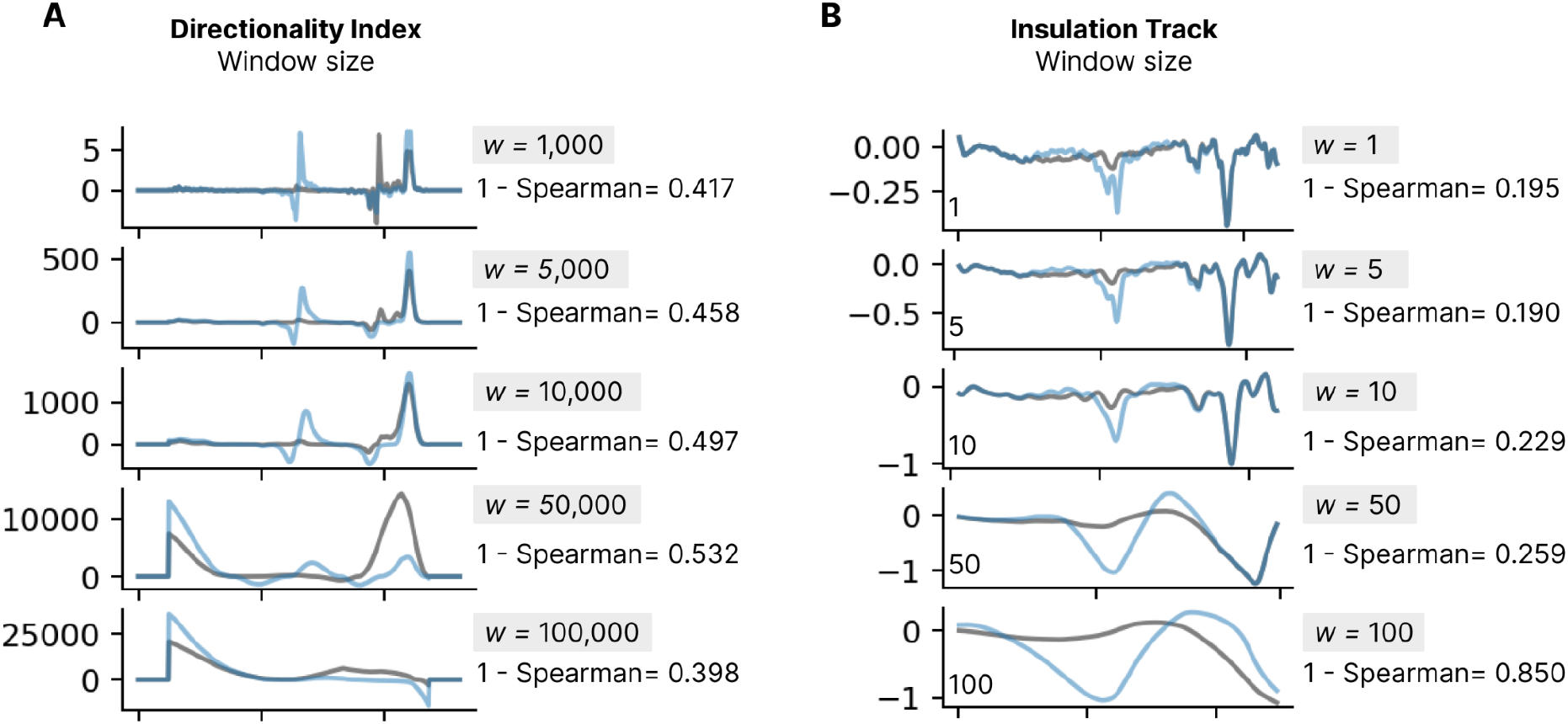
Sensitivity of directionality index and insulation tracks on parameter shifts. Directionality index (**A**) and insulation (**B**) tracks across a range of input window size choices, as well as the resulting Spearman’s correlation between the two tracks. A window size of 10 Mb was used for both approaches to produce the *in silico* scoring results in the **Results** section.

### Supplemental Tables

**Supplementary Table 1:** Method summary table.

| Method (abbreviation) | Description | Parameters | Time per 1k calls (s) |
| --- | --- | --- | --- |
| Mean squared error (MSE) | Measures the average squared difference between two flattened contact matrices. | NA | 0.24 |
| Spearman's correlation coefficient (corr) | Assesses the correlation between the intensity of corresponding pixels in two maps. | NA | 5.31 |
| Stratum-adjusted correlation coefficient (SCC) | Addresses bias due distance-dependence patterns in Hi-C data by stratifying based on their genomic distance, calculating a Pearson correlation coefficient for each stratum and aggregating the weighted stratum-specific correlation coefficients with weights derived from the generalized Cochran–Mantel–Haenszel statistic. | NA | 2.96 |
| Structural similarity index measure (SSIM) | Quantifies the perceived change in structural information of two images by incorporating three terms that correspond to structure, luminance, and contrast | NA | 2.37 |
| Eigenvector difference (eigenvector) | Calculated from the first eigenvector that corresponds to each contact frequency map. Eigenvector tracks are compared between maps using spearman's rank correlation. | NA | 41.51 |
| Directionality index difference (DI) | A measure of contact frequency bias towards either upstream sequence or downstream sequence at a locus. An inflection of DI values from negative to positive and vice versa indicates a potential chromatin boundary. | (1) size of the focal bin where upstream and downstream contact frequency is compared (2). size of upstream and downstream bins | 3.98 |
| Insulation difference (insulation) | Identifies TAD boundary-like regions by comparing the frequency of within-region contacts upstream and downstream of some point to the inter-region contacts between regions A and B | (1). size of central window (region where insulation score is calculated, x_size) (2). size of upstream/downstream windows used (a_size) | 10.98 |
| Contact decay difference (contact decay) | Measures chromatin interaction as a function of genomic distance. Interaction frequency across the contact map is ranked by genomic distance between all pairs of contact, resulting in a track of distance vs interaction frequency. As distance increases, the probability of contact between loci decreases. | (1). mean vs median | 7.17 |
| Triangle profile (triangle) | Tries to leverage our understanding that contacts are represented by different subsets of triangles within the larger map. Compares the average contact intensity within all sub-triangles of two contact maps. | NA | 4284.15 |
| TAD calling (TADs) | TAD boundaries are called by finding the local minima of the insulation profile, which is calculated using a diamond-shaped window-based method. Specifically, a square is slid along each diagonal bin of the matrix and the averaged contact frequency within each window is calculated and called as insulation score. | (1) window size (w) (2). threshold of boundary strength (3). upper bound of distance when two TAD boundaries are considered one | 8.16 |
| Loop calling (Loops) | Loops are identified by comparing with their local background. For each bin in the upper triangle window of the matrix, we first check whether it is a local maximum and then calculate the mean signals of center window surrounding the bin and also the mean signals in a donut-shape neighborhood, a lower-left neighborhood, vertical and horizontal neighborhoods around the pixel. | (1). center window size (p) (2). window size (w) (3). threshold of the ratio of center window to donut and lower-left filter (4). threshold of the ratio of center window to vertical filter (5). threshold of the ratio of center window to horizontal filter (6). upper bound of bin distance where two loops are considered as same one. | 104.18 |

### Supplemental Text

#### Basic methods

##### Mean Squared Error

The mean squared error (MSE) measures the average squared difference between two flattened contact matrices, such that

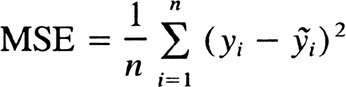

Because MSE is a measure of absolute difference, it consistently prioritizes the greatest changes in intensity between contact maps. MSE has been widely adopted across machine learning as a loss function for consistent performance and ease of use ^1–4^. Large changes between maps score highly, while visually smaller or localized changes produce lower MSE values. However, maps with differences in read count or normalization intensity will produce high MSE, despite little change in structure. For this reason, technical artifacts may dominate top map rankings scored by MSE. MSE will also deprioritize maps with large structural changes and low overall contact intensity. 2D map features will not be individually captured since the matrices are collapsed to 1D vectors.

##### Spearman’s Rank Correlation Coefficient

The Spearman’s rank correlation coefficient (ρ) assesses the correlation between the intensity of corresponding pixels in two maps by quantifying how well the relationship between the corresponding pixels can be described using a monotonic function. If the rank of intensity of all pixels in two contact maps are the same, the correlation is 1. If there is no relationship between the rank of pixel intensity between maps, the correlation is 0. Spearman’s Rank Correlation coefficient can be described as:

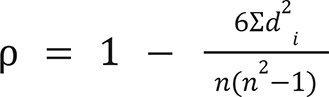

, where the number of points in the data set is represented by 𝑛, and *d*^2^ is the squared difference in the ranks of a single coordinate y_i_ between the two maps, which is summed over all points.

Correlation coefficients have been used extensively to compare contact maps^1, 4–6^. Large-scale structural changes have high scores with correlation, because the ranks of each pixel in the maps are very different. This approach works well even when the contact intensity is low because magnitude of the values is not considered when converting to rank. However, Spearman’s correlation is low even when the contact intensity is negligible (e.g. at an extreme, random noise will generate a very low correlation). The method does not pick up on small or focal changes in intensity, nor does it prioritize large-scale changes in intensity that do not change the map structure– the rank will stay the same even if the magnitude of the values change. Because matrices are flattened before calculating correlation, correlation also ignores the physical relationships between pixels of the map.

##### Stratum-adjusted Correlation Coefficient (SCC)

Contact frequency in Hi-C maps is known to exhibit a distance-dependent decay. The high similarity of the dependence pattern might bias the correlation between Hi-C maps, thus causing high, spurious correlations. SCC addresses this distance-dependence effect by stratifying Hi-C data based on genomic distance, calculating a Pearson correlation coefficient for each stratum and aggregating the weighted stratum-specific correlation coefficients with weights derived from the generalized Cochran–Mantel–Haenszel (CMH) statistic^7^. SCC values range from −1 to 1 and share a similar interpretation as standard correlations. The equation to calculate SCC can be written as:

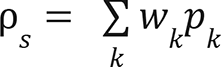

SCC was first implemented for Hi-C map comparison by Yang et al. in the R package HiCRep ^7^. By including distance-aware weights, SCC is able to measure the overall reproducibility of the Hi-C matrices better than standard correlations and is resistant to decreased resolution. However, SCC is less likely to identify small changes in TAD substructures compared to some other methods surveyed here (see **Table 1**).

##### Structural similarity index measure (SSIM)

Structural similarity index quantifies the perceived change in structural information of two images by incorporating three terms:

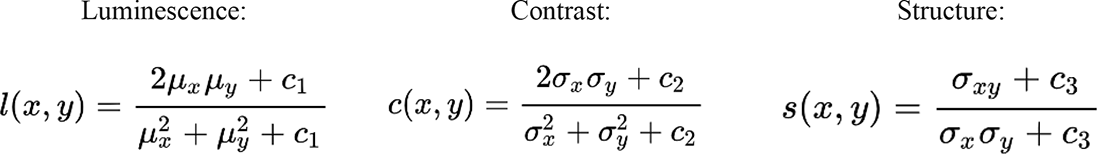

Where the integrated SSIM score is equal to:

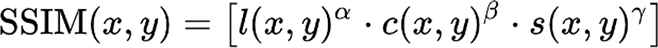

SSIM is well-suited for identifying structural changes, and unlike correlation and MSE measures, is not biased by map contrast values. For this reason it has been incorporated into Hi-C map comparison methods previously^8^. However, SSIM is sometimes very sensitive to small changes relative to larger-scale changes that may appear more pronounced to the human eye. SSIM is also sensitive to the order of the input. It should be applied to the matrix as a whole (not vector-by-vector) as it is designed to account for neighboring values. NaN values must be interpolated or masked to zero.

#### Map-informed methods

##### Eigenvector difference

This method is inspired by genomic compartments, which are called by calculating the first eigenvector from Hi-C contact maps and assigning each genomic region to its sign ^6^. Similarly, eigenvector difference is calculated from the first eigenvector that corresponds to each contact frequency map, creating a vector annotated at each bin for both maps. These vectors are then compared using spearman’s rank correlation. Because the components can have different signs that are arbitrarily assigned, MSE is not used for this method as it is sensitive to these signs and would result in falsely high scores when the maps are assigned opposite signs.

##### Directionality Index (DI)

The Directionality Index (DI) is a measure of contact frequency bias towards either upstream sequence or downstream sequence at some DNA locus. An inflection of DI values from negative to positive and vice versa indicates a potential chromatin boundary, where DI can be calculated by:

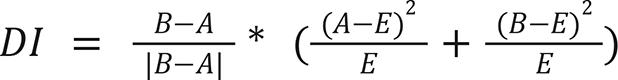

Where A is the number of reads (or average normalized frequency value) that map from a given locus to upstream bins, B is that value for downstream bins, and E is the expectation under the null hypothesis, equal to 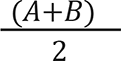.

DI was first proposed by Dixon et al. in 2012 ^9^. It depends on two parameters: the size of the focal bin whose relative upstream and downstream contact frequency is being compared, and the size of the upstream and downstream bins (40 kb and 2 Mb in the original publication). To create a composite DI disruption score for a variant, DI is calculated for each locus in the region of both maps and compared using MSE or correlation. This composite score is subject to the caveats of the chosen comparison method.

##### Insulation score

Also known as the ratio score or boundary score, this method seeks to identify TAD boundary-like regions by comparing the frequency of within-region contacts upstream (A) and downstream (B) of some point X to the inter-region contacts between regions A and B ^10, 11^. The higher the ratio is in a given region, the more likely this region is to be a TAD boundary. The metric is calculated as follows:

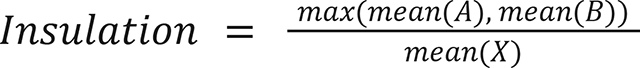

, where X quantifies the frequency of local contacts within the central region spanning 20 kilobases, and A and B represent contact frequency in the regions upstream and downstream of X, respectively, spanning 200 kilobases each.

Correlation or MSE can be applied to the insulation tracks of two conditions for a scalar disruption score. The magnitude of the insulation score is dependent on differences in contact intensity between the two maps at each bin, and therefore is sensitive to global change in contact frequency. Variants in regions of DNA with wider ranges of contact intensity (high contrast) may have inflated insulation scores relative to other regions. This method can potentially be improved by adjusting the following parameters: the size of central window X, i.e. the region for which the insulation score is being calculated (default: 20kb), and the size of upstream/downstream windows A and B (default: 200kb)

##### Contact probability decay

Contact decay, or the P(S) curve, measures chromatin interaction as a function of genomic distance ^12, 13^. Interaction frequency across the contact map is ranked by genomic distance between all pairs of contact, resulting in a track of distance vs interaction frequency. As distance increases, the probability of contact between loci decreases. Decay curves may be calculated at a given resolution such that the chromosome is divided into n = L/r bins, where L is the chromosome length and r is resolution. Across an *n* x *n* contact map, the contact frequency of each entry A_i,j_ is ordered by the distance between loci, i-j. A steeper decay in contact frequency indicates a greater distance between further loci, while a shallow contact decay suggests more interaction between distant loci. Contact decay measures a global signal of relative interaction increase or decrease, but will not be sensitive to local structural changes to contact matrices.

##### Triangle Method

Basic methods (correlations, MSE) all ignore the physical relationships between pixels of the map when they are flattened into vectors. They are therefore over-simplified characterizations of the relationships between maps. This method tries to leverage our understanding that contacts are represented by different subsets of triangles within the larger map to address this gap. The triangle-based method compares the average contact intensity within all sub-triangles of two contact maps. In comparison to MSE and correlations, the flattened representation of the map is the average contact intensity of all sub-triangles instead of just each pixel on its own. The flattened representations can then be compared with either MSE or a correlation method.

The performance of this method depends on the correlation or MSE used over the sub-triangles (see their individual pros and cons). The advantage over those basic methods is that triangle comparison is more feature-informed to capture relevant contact relationships. Because there are so many more smaller triangles than larger triangles, this method likely prioritizes more local changes; however, one could weight the triangles or subset to only the larger or smaller sub-triangles to prioritize only larger or smaller scale interactions. One caveat is that this method is significantly slower than other methods, but speed can be improved by creating lower resolution maps before computing.

#### Feature-informed methods

##### TADs

TAD boundaries are called by finding the local minima of the insulation profile, which is calculated using a diamond-shaped window-based method proposed by Crane et al. ^10^. Specifically, a square (a *W* x *W* diamond-shaped window) is slid along each diagonal bin of the matrix and the averaged contact frequency within each window is calculated and called as insulation score. Bins with a low insulation score indicate a high insulatory effect, thus the bins reaching the local minima are identified as candidate TAD boundaries. The boundary strength is calculated for each local minima using peak prominence and candidates with strength above a threshold are referred to as TAD boundaries. The scores for the bins at the end of the diagonal and within the window size are not calculated. The overlap, gain, and loss of TAD boundaries between two Hi-C matrices are reported to show their consistency and changes. The boundary locations within a set resolution *r* are considered the same. This method could be further improved by changing the following parameters: window size (*w*), threshold of boundary strength, and upper bound of distance when two TAD boundaries are considered the one.

##### Loops

Chromatin loops are the positions where a pair of loci showing closer proximity compared to loci lying between them, corresponding to pixels with higher contact frequency than the ones in their neighborhood. We identify loops by comparing regions with their local background, as in HiCCUPS^14^. Specifically, for each bin in the upper triangle window of the matrix, we first check whether it is a local maximum (across neighborhood window size *w*) and then calculate the mean signal of center window (window size *p*) surrounding the bin as well as the mean signal in a donut-shape neighborhood, a lower-left neighborhood, vertical and horizontal neighborhoods around the pixel. The bins enriched above its neighborhood with ratios of mean signals of the center window to the neighborhoods higher than certain thresholds are considered as candidate loops. The bins at the corners are not considered. Loops that are the same, gained, lost between two Hi-C matrices are identified. The loops that are located within a window of size *r* of one another are treated as the same. This method can potentially be improved by adjusting the following parameters: the center window size (*p*), window size (*w*), threshold of the ratio of center window to donut and lower-left filter, threshold of the ratio of center window to vertical filter, threshold of the ratio of center window to horizontal filter, and the upper bound of bin distance where two loops are considered as same one.

